# A reciprocal inhibition model of alternations between non-dissociative and dissociative states in patients with PTSD

**DOI:** 10.1101/795732

**Authors:** Toshinori Chiba, Kentaro Ide, Shuken Boku, Jessica E. Taylor, Hiroyuki Toda, Tetsufumi Kanazawa, Sumie Kato, Yuka Horiuchi, Akitoyo Hishimoto, Toru Maruyama, Taisuke Yamamoto, Miyako Shirakawa, Ichiro Sora, Mitsuo Kawato, Ai Koizumi

## Abstract

**Objective:** Traumatic life-events can leave individuals with contrasting posttraumatic stress disorder (PTSD) symptoms, including re-experiencing and avoidance. Notably, patients with PTSD are known to periodically switch between two opposing attentional biases; namely, toward threat and away from threat. We hypothesized that reciprocal inhibition between the amygdala and ventromedial prefrontal cortex (vmPFC) may induce alternations between these attentional biases, which in turn may contribute to the re-experiencing and avoidance symptoms, respectively.

**Methods:** To test this reciprocal inhibition model, we performed an experiment to measure the attentional biases of patients with PTSD. We examined the differential relationships between PTSD symptom clusters (re-experiencing/avoidance) and attentional biases (toward/away from threat). Additionally, we performed a meta-regression analysis to examine the role of amygdala reactivity in the imbalance between re-experiencing and avoidance symptoms.

**Results:** We found that attentional bias toward threat was selectively associated with re-experiencing symptoms whereas attentional bias away from threat was selectively associated with avoidance symptoms. Meta-regression analysis based on twelve participant populations (total N = 316) revealed that left amygdala activity was positively correlated with the severity of re-experiencing symptoms relative to avoidance symptoms.

**Conclusions:** Our findings support the hypothesis that reciprocal inhibition of common neural circuits may underlie the switch between attentional biases toward and away from threat as well as that between re-experiencing and avoidance symptoms. Re-experiencing and avoidance/emotional numbing are the core symptoms used to distinguish between the non-dissociative and dissociative PTSD subtypes. The reciprocal inhibition mechanism may help elucidate the mechanisms underlying those PTSD subtypes.

## INTRODUCTION

Posttraumatic stress disorder (PTSD) is a debilitating disorder that develops after experiencing life-threatening traumatic events. It is a paradoxical disorder because individuals with PTSD can display seemingly opposing symptoms. Specifically, they may display re-experiencing symptoms where they automatically engage with traumatic cues but may also display avoidance/dissociative symptoms where they actively stay away from such cues (1, 2).

Patients with strong dissociative symptoms are categorized as having the dissociative PTSD subtype, which was added to the fifth edition of the Diagnostic and Statistical Manual of Mental Disorders (DSM-V) (1). These participants exhibit multiple characteristics, including lower treatment response and a higher suicidal risk (3), which are distinct from those exhibited by typical patients with PTSD (the “non-dissociative subtype”). The dissociative subtype is characterized by hypoarousal (avoidance/dissociation) symptoms and was postulated to be driven by over-inhibition of the amygdala by the ventromedial prefrontal cortex (vmPFC) (1, 2). Conversely, the non-dissociative subtype is characterized by hyperarousal (re-experiencing/hypervigilance) symptoms and was postulated to be driven by under-inhibition of the amygdala by the vmPFC (1, 2).

It remains unclear whether patients remain stable or spontaneously fluctuate between either the non-dissociative or dissociative state. The current theoretical and clinical consensus is that PTSD is a dynamic disorder that involves fluctuations between the non-dissociative and dissociative states (1, 4–6). In line with this, previous studies reported fluctuating behavioral, physiological, and neuroimaging responses within individual patients with PTSD (4–7), which possibly reflects variability of their inner dissociative state levels. However, it is unclear whether the observed fluctuations in responses specifically reflect switches between the non-dissociable and dissociable states or if they just reflect random jitters in responses within one stable state. Dissociating these two possibilities would benefit from a comprehensive framework encompassing both within- and between-patient fluctuations in their states.

One effective way to assess within- and between-patient alternations in non-dissociative and dissociative states may be to examine the fluctuations in attentional bias within individual patients (i.e., toward or away from threat) (7, 8). Attentional bias toward threat allows one to detect imminent threats rapidly and to adaptively avoid them (9). While bias toward threat is found to be overemphasized in most anxiety disorders, including PTSD (10, 11), some patients with PTSD have been reported to instead display attentional bias away from threat (12). The heterogeneity of these results suggest that attentional biases are unstable and fluctuate over time in patients with PTSD. This attentional fluctuation is usually viewed as a reflection of “instability” in threat monitoring in patients with PTSD (7, 8). In traditional analyses of attentional bias, an overall bias toward threat is associated with non-dissociative characteristics (11) while an overall bias away from threat is associated with dissociative characteristics (13, 14). Although researchers have not determined the mechanism involved in this “instability” of attentional bias, one possibility is that such “instability” may emerge from the same mechanism as that causing within-patient alternation between the non-dissociative and dissociative states. We tested if a single neural network model could comprehensively explain the fluctuations in both attentional bias and dissociative states.

Fluctuations between the non-dissociative and dissociative states in PTSD may be explained by neural circuits that can produce rhythmic fluctuations between two states rather than by those that only possess stable equilibrium points. Generally, reciprocally inhibiting circuits produce such rhythmic fluctuations (15–17). Here, we propose a conceptual neural network model to explain the two opposing PTSD states and attentional biases. To test the predictions of this model, we conducted a behavioral experiment to measure attentional bias among patients with PTSD. To further test the neural-level assumptions underlying the model, we performed a meta-regression analysis of findings from previous neuroimaging studies on the association between amygdala reactivity and non-dissociative/dissociative symptom clusters using data from 12 populations (total N = 316).

### Reciprocal inhibition model: proposal for attention oscillations and alternations of dissociative/non-dissociative states

In reciprocal inhibition, two distinct neural circuits alternatively dominate each other via mechanisms such as post-inhibitory rebound and spike frequency adaptation (17, 18). For example, neurons in one circuit that are initially inhibited may escape from the inhibitory modulation when their intrinsic membrane properties allow them to cross the spike threshold, which then inhibits neurons in the initially inhibiting circuit. Thus, two competing neural circuits take turns to induce two alternative states.

We hypothesized that reciprocal inhibition between the amygdala and vmPFC may contribute to alternations between the non-dissociative and dissociative states in PTSD (Figure 1A). When the amygdala is activated, the vmPFC is suppressed, causing attention to be biased toward threat. This subsequently induces re-experiencing symptoms (Figure 1A left). Conversely, when the vmPFC is activated, the amygdala is suppressed, causing attention to be biased away from threat. This subsequently induces avoidance/dissociative symptoms (Figure 1A right).

**Figure 1.**
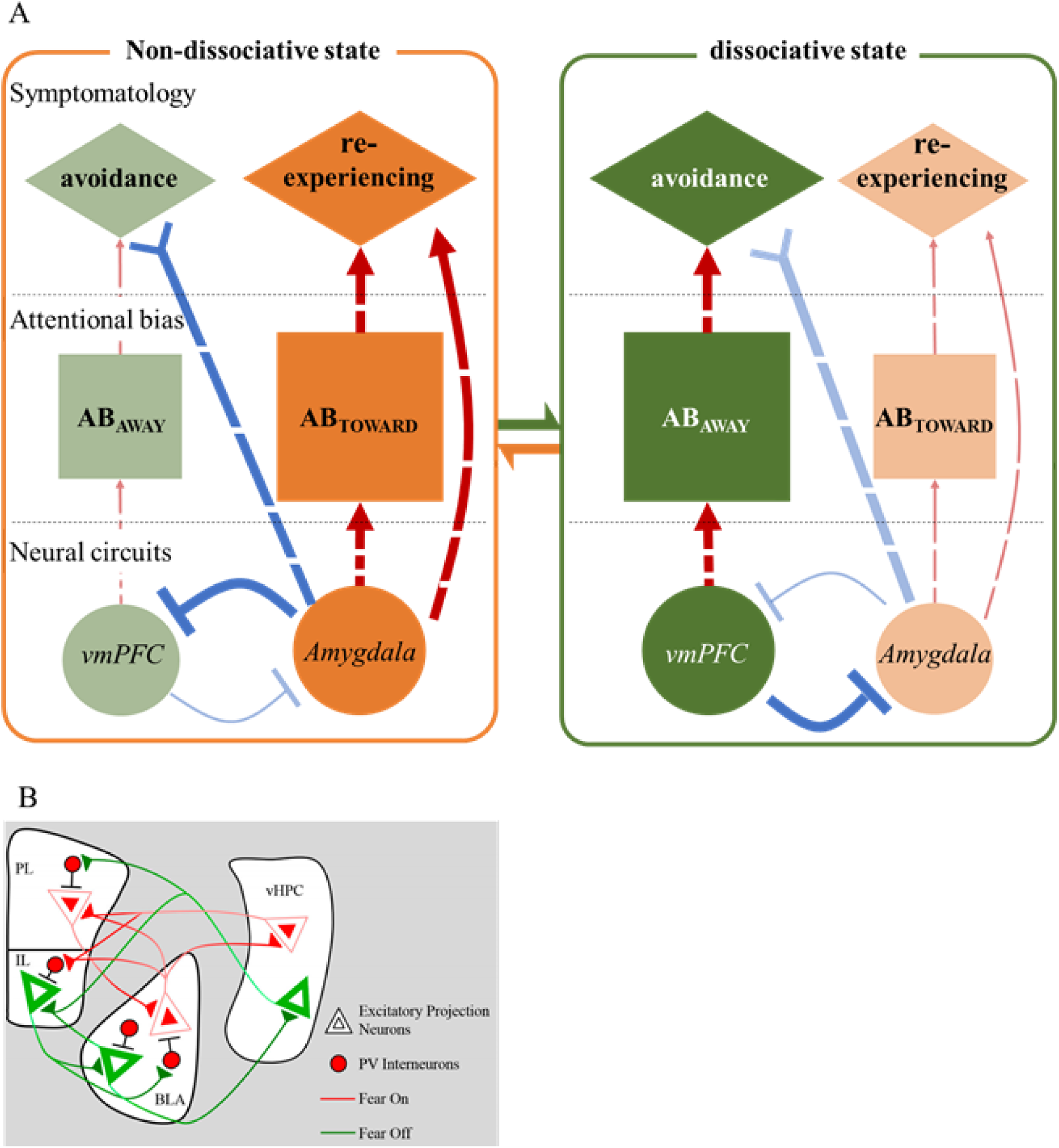
The reciprocal inhibition model, which explains fluctuations between the dissociative and non-dissociative states of PTSD. (A) In the reciprocal inhibition model, reciprocal inhibition between the amygdala and the vmPFC would contribute to alternations between the non-dissociative and dissociative states in PTSD. Activation of the amygdala causes attention to be biased towards threat, subsequently causing re-experiencing symptoms to manifest (Figure 1 A left). Conversely, activation of the vmPFC causes attention to be biased away from threat, subsequently causing avoidance/dissociative symptoms to manifest (Figure 1 A right). (B) The microcircuits that might underlie reciprocal inhibition. The amygdala (BLA “fear -on” neurons) and the vmPFC (IL “fear-off” neurons) reciprocally inhibit each other via activation of GABAergic inhibitory interneurons. PL: prelimbic (medial prefrontal) cortex, IL: infralimbic (medial prefrontal) cortex, vHPC: ventral hippocampus, BLA: basolateral amygdala, PV: parvalbumin Figure 1 (B) is adopted from Zimmermann et. al. with no permission yet asked (45).

In a neural circuit model with only a stable equilibrium point, re-experiencing and avoidance symptoms could have stable associations with attentional bias and neural reactivity over time. In contrast, our model predicts that these two distinct symptom clusters are distinctly associated with these measures, depending on the predominant state. The activity of the amygdala averaged over a time-window is predicted to be composed of the average of high and low activities from the amount of time spent in the non-dissociative and dissociative states, respectively. Amygdala activity should be positively correlated with re-experiencing symptoms during the non-dissociative state and negatively correlated with avoidance symptoms during the dissociative state. Therefore, overall amygdala activity is predicted to be positively correlated with subtraction of avoidance scores from re-experiencing scores.

The proposed model is circumstantially supported by the following empirical data. The amygdala and vmPFC appear to be generally reciprocally inhibitorily connected. A previous resting-state functional connectivity analysis of healthy individuals demonstrated that spontaneous fluctuations in the blood oxygen level-dependent (BOLD) signals of an amygdala subdivision were negatively correlated with those in the vmPFC (19). Moreover, during fear-processing, the BOLD signal of the amygdala is negatively correlated with that of the vmPFC in healthy adults (20–26). In patients with PTSD, abnormality of the amygdala-vmPFC network is widely observed (27–35). The amygdala-vmPFC network is characterized by a pattern of predominant bottom-up and top-down connectivity in the non-dissociative and dissociative subtypes, respectively (36). Additionally, amygdala activity is positively correlated with attentional bias toward threat (37–39) and re-experiencing symptoms (11, 40, 41) while vmPFC activity is positively correlated with attentional bias away from threat (14, 42) and avoidance/dissociative symptoms (36, 43, 44). Findings from rodent studies indicate precise microcircuits that could at least in part underlie the reciprocal inhibition proposed by our model. In these studies, reciprocal inhibition is shown to occur between the amygdala “fear-on” neurons and vmPFC “fear-off” neurons via GABAergic inhibitory interneurons (Figure 1B) (25, 45–51).

Here, in our meta-analysis we were able to analyze reactivity in the amygdala but not in the vmPFC. This is because most previous studies did not focus on the vmPFC but focused instead on related, overlapping, or sub-regions of the vmPFC (52–54). Therefore, there was an insufficient number of previous studies with coherent definitions of the vmPFC for meta-analysis.

Unlike other dissociative symptoms, such as depersonalization or derealization, avoidance symptoms are a necessary requirement to meet the full PTSD criteria; therefore, they are found in all patients with PTSD (1). This allowed us to use scores from the avoidance symptom cluster of the DSM-IV scores to index the severity of dissociative symptoms in each of our patients. The dissociative PTSD subtype was first officially defined in the DSM-V as being characterized by depersonalization and derealization symptoms. Emotional numbing symptoms, which are the main type measured in the DSM-IV avoidance symptom cluster and are also thought to represent a dissociative state, were not included in this official definition. As further elaborated in the Discussion section, this suggests that the dissociative/non-dissociative PTSD subtypes are related but do not directly correspond to dissociative/non-dissociative states.

## METHODS

### Behavioral analysis

### Participants

We enrolled 20 patients with PTSD (2 males, 18 females; mean age = 41.2 years; range = 22-53 years) from the Flower of Light Clinic for Mind and Body (n = 15) and the Chiyoda-Shinryou clinic (n = 2), which are both located in Tokyo, and the Shinchi-clinic (n = 3) located in Osaka. All the participants were diagnosed with DSM-IV PTSD resulting from domestic violence (n = 5), childhood abuse (n = 2), an unpleasant sexual experience (n = 2), or a combination of these (n = 11) according to the Clinician-Administered PTSD Scale (CAPS: mean score = 80.6, SD = 20.2, range = 51-119, see Supplementary Table 1 for details). These participants reported strong fear when viewing pictures of angry male faces; were not taking psychotropic medication; had not suffered traumatic brain injury or loss of consciousness; and did not have any lifetime history of psychosis, alcohol abuse, or substance abuse. This study was approved by the Ethics Committees of the Central hospital of National Defense Force and Advanced Telecommunications Research Institute International (ATR). All the participants provided written informed consent.

#### Experimental tasks and procedures

The breaking continuous flash suppression (b-CFS) task was adapted from the study by Yang, Zald, and Blake (55). CFS renders a target stimulus invisible by presenting it to one eye while presenting a mosaic pattern to the other eye (56). The b-CFS task assesses the detection time of stimuli masked by binocular suppression (Figure 2A). Grayscale pictures of six male faces were obtained from the ATR Facial Expression Image database (DB99) and used as target stimuli. These pictures depicted angry or neutral expressions and were cropped in a circular shape to include the brows, eyes, nose, and mouth. All pictures were equated for contrast and luminance. Visual stimuli were presented using MATLAB (MathWorks, Inc.) with the Psychophysics Toolbox extensions (57, 58). Stimuli were presented dichoptically through an Oculus Rift head-mounted display. To facilitate binocular fusion, one black “fusion frame” was displayed to each eye. A black fixation cross was drawn in the center of each fusion frame and the participants were instructed to remain fixated. Target stimuli were presented covering 1 of 4 quadrants within the fusion frames. Twelve pictures (6 males × 2 expressions) were presented once in every quadrant in a randomized order resulting in a total of 48 trials. To render stimuli invisible, Mondrian-like masks were flashed at 10 Hz to the dominant eye while target stimuli were presented to the other eye.

**Figure 2.**
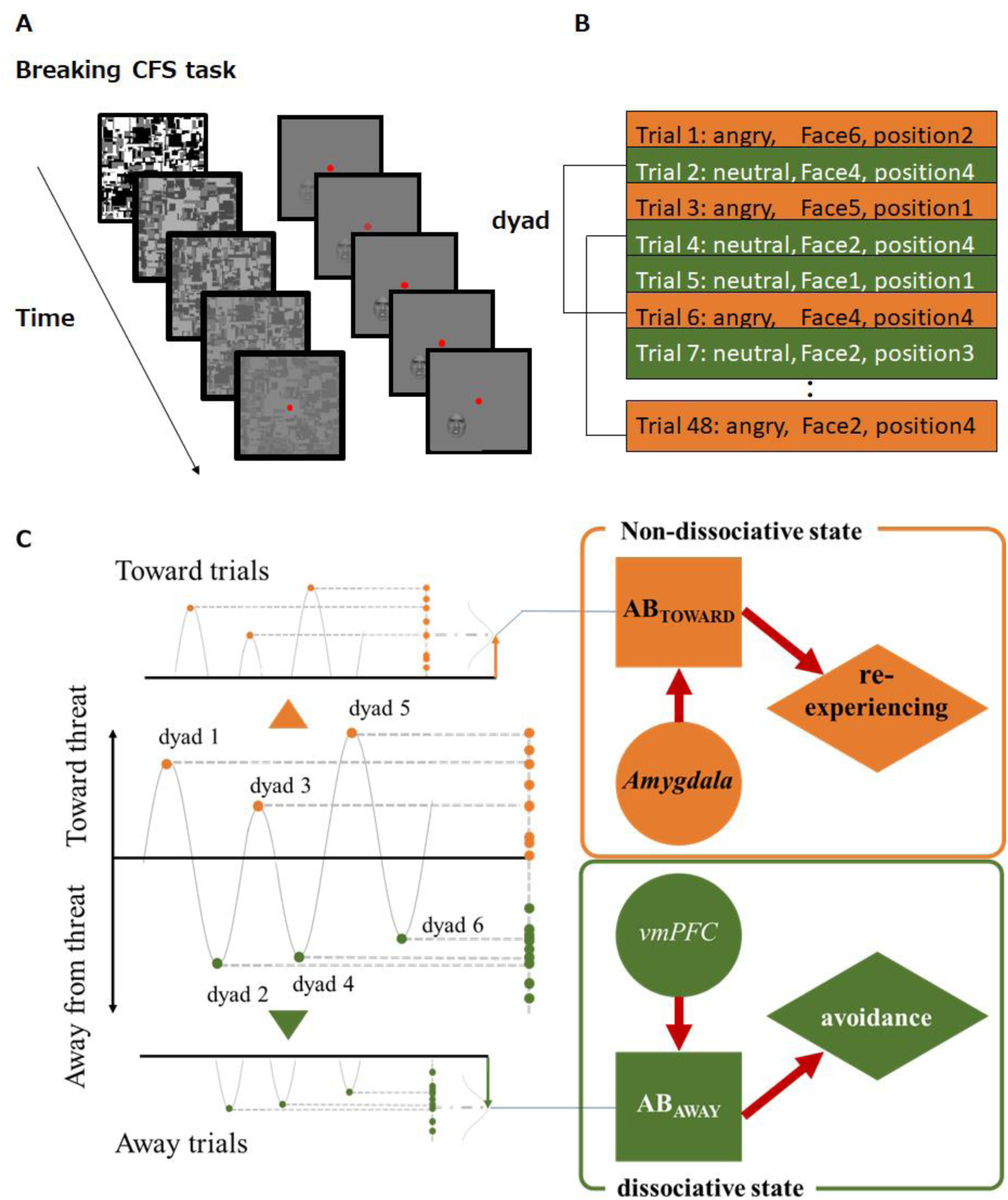
Examples of CFS trial presentation and the putative generation of attentional bias during non-dissociative and dissociative states. (A) An example of a breaking continuous flash suppression (b-CFS) trial presentation. During each trial, 1 of 6 different faces (Face1-Face6; angry or neutral expressions) was presented within 1 of 4 quadrants (position1 to position4) on a frame presented to one eye. (B) Each dyad consisted of one neutral face and one angry face trial with consistent face identity (e.g. Face1) and position (e.g. position1). (C) For each dyad, the attentional bias score was defined as the difference between the reaction time to the angry face and that to the neutral face. Positive and negative values represent attentional bias toward and away from threat, respectively. We averaged attentional bias scores of all the positive and negative dyads separately and defined them as AB_TOWARD_ and AB_AWAY_. The standard deviations (SDs) of all of the dyads in total (not split by valance) are comparable with attention-bias variability (Var) measures used in previous studies. The averages of all the dyads in total are comparable with traditional attentional bias (TAB) measures used in previous studies. Since the biases toward and away from threat are indicative of re-experiencing and avoidance, respectively, we conceptually illustrated the putative generation of attentional bias during non-dissociative and dissociative states. That is, AB_TOWARD_ and AB_AWAY_ are associated with re-experiencing and avoidance states, respectively.

Each trial started with a fixation period of 4 s. Subsequently, suppression masks were flashed to the dominant eye, and the target stimulus was gradually faded into the non-dominant eye by linearly increasing its contrast over 1 s. During the 1-7 s period after trial onset, the contrast of the CFS masks was slowly decreased to zero. Stimuli were displayed until the participants pressed a key to indicate the quadrant in which the target stimulus (or any part of the target stimulus) emerged from suppression (55, 59). The participants were required to respond as quickly as possible without compromising accuracy. The participants’ response cleared the screen.

To familiarize the participants with the procedure, they were presented with 12 practice trials (each of the 12 target stimuli was presented once) at the beginning of the experiment. Before beginning the experiment, eye dominance was examined using the hole-in-a-card test (60).

#### Two attentional bias scores

Reaction times for correct trials were the main outcome measured in this study. Trials with incorrect responses and one extreme outlier trial, in which the response time was more than five standard deviations above the participants’ mean for that particular condition, were excluded from subsequent analysis (<3% of all trials). There was no difference in the percentage error rate between stimulus facial expressions (i.e., fearful or neutral) (t = 1.228, df = 19, *p* = 0.23). During each trial, 1 of the 6 different faces (Face1 to Face6; angry or neutral expressions) was presented within 1 of 4 quadrants (position1 to position4) on a frame presented to one eye. We refined previously used attentional bias parameters for each participant (61) to generate a more stable index that was less influenced by the effects of face identity and the presented position. To this end, stimuli were paired in dyads consisting of one neutral face trial and one angry face trial with consistent face identity (e.g., Face1) and position (e.g., position1). Next, for each dyad, the attentional bias score was defined as the difference between the reaction time to the angry face and the reaction time to the neutral face. Since positive attentional bias scores represent attentional bias toward threat, we averaged all of the positive attentional bias scores and defined them as attentional bias score toward (AB_TOWARD_). Similarly, since negative attentional bias scores reflect attentional bias away from threat, we averaged all the negative attentional bias scores and defined them as AB_AWAY_. The absolute value of the participants’ AB_AWAY_ was used in subsequent analyses to allow easy comparisons of the magnitudes of attentional bias scores.

#### Statistical analysis of attentional bias scores and PTSD symptoms

A stepwise multiple regression analysis was performed to evaluate the relationships between three symptom clusters (re-experience, avoidance, and hypervigilance) and each attentional bias score (AB_TOWARD_ and AB_AWAY_). In this regression, an automatic statistical model selection procedure was adopted as an exploratory means of identifying which symptom cluster/combination best explains the attentional bias scores. To test if the current data met the assumption of multicollinearity, we computed the intercorrelation between the predictor variables (re-experience, avoidance, and hypervigilance). In all analyses, p < 0.05 was considered statistically significant.

### Meta-regression analysis of the relationship between amygdala fMRI signals and PTSD subcluster scores

#### Systematic literature search

We performed a systematic literature search using PubMed between April 1, 2019 and April 20, 2019. The following keywords were used in our search: “fMRI” OR “functional magnetic resonance imaging” combined with AND “PTSD” OR “posttraumatic stress disorder” OR “acute stress disorder”. The present systematic review follows the Preferred Reporting Items for Systematic Reviews and Meta-Analyses (PRISMA) guidelines. The inclusion criteria are presented in the PRISMA flow chart (Supplementary Figure 1). Neuroimaging studies were included if they (1) included traumatized individuals as participants; (2) were published in English; (3) compared fMRI BOLD signals to (a) threatening stimuli vs. neutral stimuli, (b) stimuli that included threatening/unpleasant stimuli vs. stimuli that did not include threatening/unpleasant stimuli, or (c) threatening/unpleasant stimuli vs. baseline; and (4) reported (a) PTSD subcluster scores and (b) amygdala BOLD signals as z-scores or t-stats. Titles and abstracts were screened for eligibility by one assessor (TC; screening phase, n = 323). The full texts of all the finally included studies were examined in detail and independently selected by two assessors (KI, TC; n = 191). All the reference lists of the reviewed papers were examined to identify other eligible studies.

#### Statistical analysis of amygdala reactivity and PTSD symptoms

For participants from each study selected for the meta-regression analysis, PTSD symptom imbalance was defined as the difference between the patients’ avoidance scores and their re-experiencing scores. Before subtraction, symptom scores were normalized so that the highest possible value was one and the lowest possible value was zero. Different studies measured their patients’ levels of PTSD using different versions of the DSM. The variables that contribute to avoidance scores in the DSM-IV protocol are divided in that some contribute to the avoidance scores and others to the emotional numbing scores in the DSM-V. Therefore, in studies where the DSM-V was used, the participants’ avoidance and emotional numbing scores were summed before being normalized to allow for direct comparisons between these studies and those that used the DSM-IV. Pearson correlation values were calculated between symptom imbalance and z-scores that represented left and right amygdala activity separately. If the study did not provide a z-score, the t-stat was transformed into a z-score using the SPM12 built-in-function.

## RESULTS

### Correlation between attentional bias scores and PTSD symptoms

We did not find multicollinearity in the data. The coefficients for the correlations between re-experiencing and avoidance (r = 0.69), avoidance and hypervigilance (r = 0.53), and re-experiencing and hypervigilance (r = 0.53) were below the suggested cut-off value of 0.8 (62). The percentage of “toward trials” (Figure 2C) was 46.3% (range 39.1-58.3%), which indicates that all the patients alternated between toward and away trials and is consistent with a previous study that showed alternations between the two attentional states (61). This is also consistent with another previous study in which patients with PTSD showed a mixture of hyperarousal and dissociative responses within the same experiment, indicating that multiple alternations between the non-dissociative and dissociative states may occur within a time range of 5-10 minutes (63). A stepwise regression analysis revealed that the re-experiencing symptom cluster was a significant predictor of AB_TOWARD_ (overall model: r^2^ = 0.20, df = 1, 18, *p* = 0.0499; re-experiencing: beta = 0.44; Figure 3A left), whereas the avoidance symptom cluster was a significant predictor for AB_AWAY_ (overall model: r^2^ = 0.22, df = 1,18, *p* = 0.036; avoidance: beta = 0.47; Figure 3A right). The order of variables added into this model was automatically determined by the algorithm and was based solely on the t-statistics of their estimated coefficients. A higher re-experiencing score predicted a greater AB_TOWARD_ (Figure 3B) whereas a higher avoidance score predicted a greater AB_AWAY_ (Figure 3C) (see Supplementary Table 1 for details). Interestingly, a similar relationship was observed between the PTSD symptom clusters and the variability within each direction of attention bias (AB_TOWARD_ and AB_AWAY_). Specifically, the re-experiencing score was significantly correlated with the variability within AB_TOWARD_ (r^2^ = 0.31, df = 1,18, *p* = 0.01) whereas the avoidance score was significantly correlated with the variability within AB_AWAY_ (r^2^ = 0.37, df = 1,18, *p* < 0.01).

**Figure 3.**
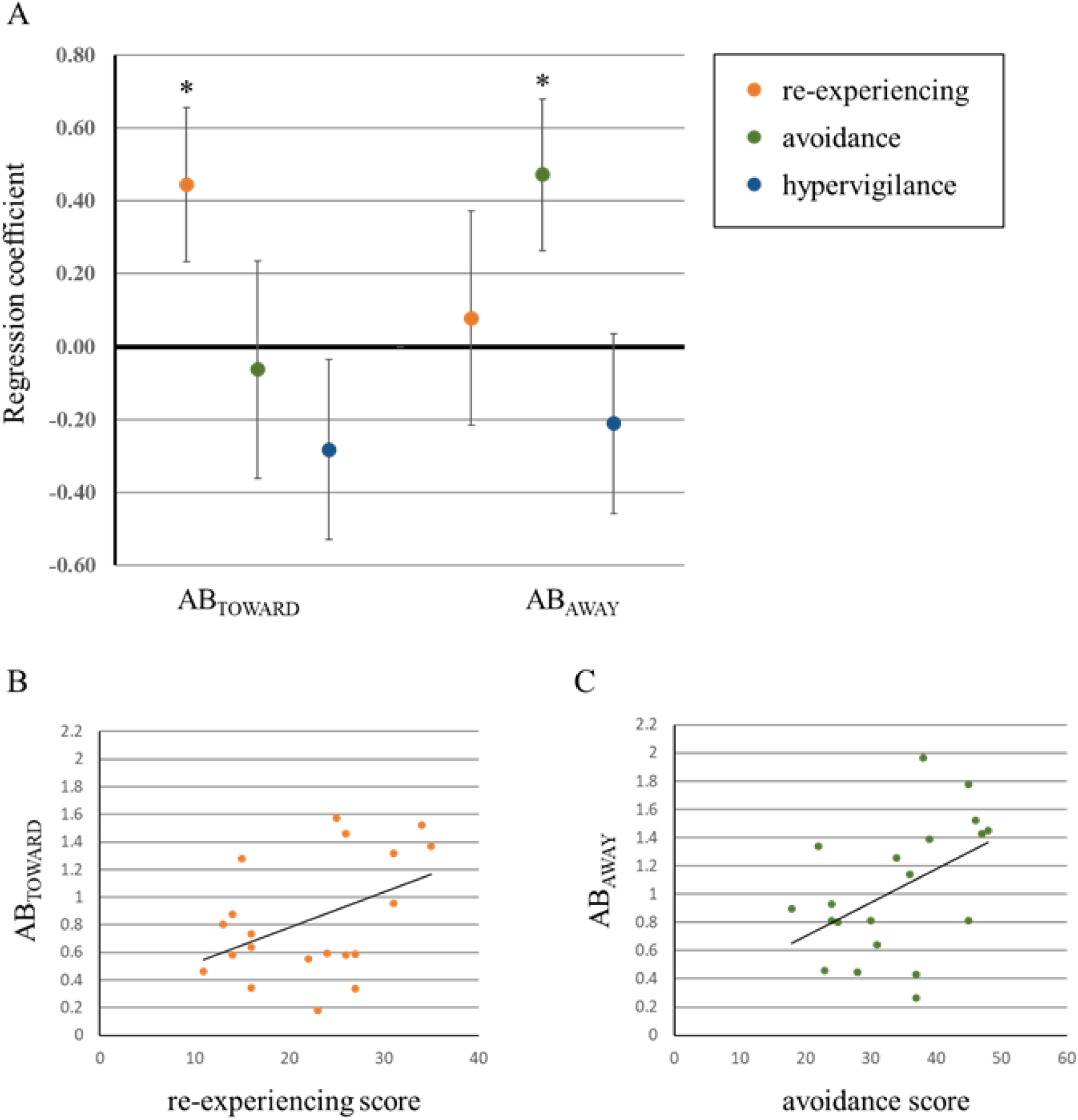
PTSD symptom clusters and attentional bias scores: results of the stepwise regression analysis. (A) AB_TOWARD_ was associated with re-experiencing, and AB_AWAY_ was associated with avoidance. For demonstrative purposes, results of single-variable regression are shown here to demonstrate the relationships between re-experiencing and AB_TOWARD_ (B) and avoidance and AB_AWAY_ (C). * *p* < 0.05

### Relationship between amygdala fMRI signals and symptom imbalance: results from the meta-regression analysis

The data from 12 participant populations (total N = 316) were extracted from 9 studies for the meta-regression analysis. Among these, left amygdala activity was reported in 8 populations while right amygdala activity was reported in 11 populations. Symptom imbalance was significantly correlated with left amygdala activity (r = 0.70, *p* = 0.028, one-tailed: Figure 4A) but not with right amygdala activity (r = 0.15, *p* = 0.66, one-tailed: Figure 4B).

**Figure 4.**
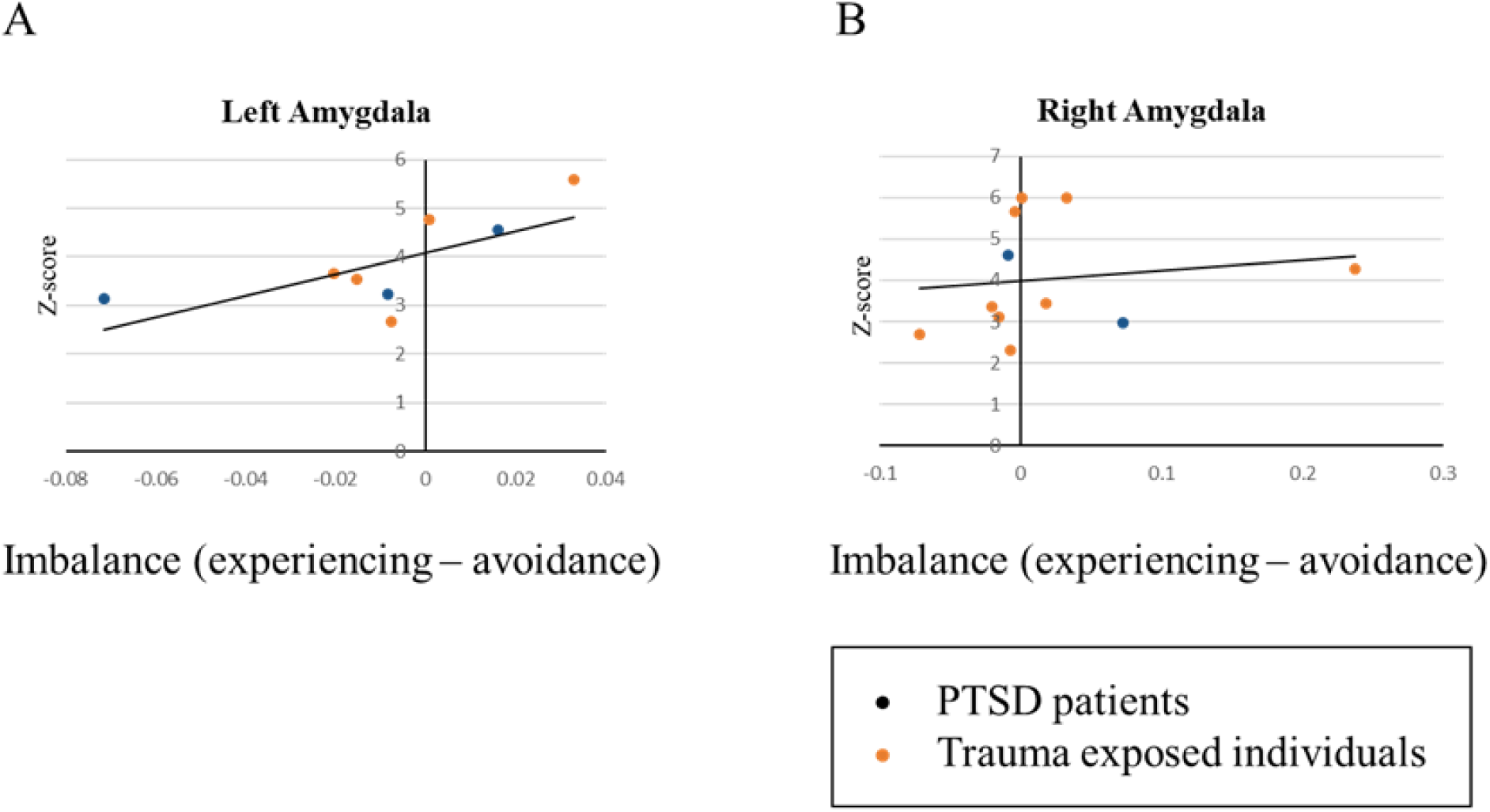
PTSD symptom imbalance and amygdala BOLD response: Results of the meta-regression analysis. Each dot represents the mean value for one participant population from systematically selected studies (Supplementary Figure 1, Supplementary Table 2). The x-axis denotes the PTSD symptom imbalance, which was defined as the difference between avoidance and re-experiencing scores. Before subtraction, symptom scores were normalized so that the highest possible value was one and the lowest possible value was zero. The y-axis denotes the fMRI BOLD signal to threat, expressed as z-scores. Left, but not right, amygdala BOLD signal was positively correlated with symptom imbalance (left amygdala: r = 0.70, *p* = 0.028, one-tailed, right amygdala: r = 0.15, *p* = 0.66, one-tailed).

## DISCUSSION

Although studies have reported an array of clinically vital differences between patients with dissociative and non-dissociative PTSD (1, 36, 64), little is known about whether individual patients experience fluctuations between the dissociative and non-dissociative states. In this study, we hypothesized that reciprocal inhibition between the amygdala and vmPFC might generate rhythmic oscillations between these states within individual patients. Our behavioral experiment and meta-regression analysis of previous studies provided supportive data for this model at two distinct levels. At the between-individual level, re-experiencing symptoms were selectively correlated with a higher attentional bias toward threat (AB_TOWARD_), while the avoidance symptoms were selectively correlated with a higher attentional bias away from threat (AB_AWAY_). This is consistent with the prediction that attentional bias fluctuations synchronize with alternations between the dissociative and non-dissociative states, rather than simply reflecting random fluctuations in brain states. The meta-regression analysis revealed that the imbalance between re-experiencing and avoidance symptoms was correlated with left amygdala reactivity to threat. This result is consistent with the hypothesis that symptom fluctuations are produced from reciprocal inhibition between the amygdala and vmPFC. Overall, we provided empirical data at the between-individual and meta-regression analysis levels that are consistent with the reciprocal inhibition model.

Our results also suggest that the findings obtained from subtype-wise analyses may be relevant at the individual patient level. For example, we can hypothesize that treatment response could be predicted by the imbalance between amygdala and vmPFC activity within individual patients. This is because patients with the dissociative PTSD subtype usually show a lower treatment response (65, 66). Further, exaggerated amygdala reactivity, which is a typical characteristic of the non-dissociative subtype, predicts a poor response to exposure-based therapy (67). An individual patient may show a smaller or greater response to an administered treatment when the amygdala/vmPFC imbalance is large or small, respectively. This is especially important since one reciprocal inhibitory circuit can produce a whole range of different oscillation frequencies under the influence of different neuromodulators (17). Reciprocal inhibition between the amygdala and vmPFC may not only induce a switch of states within the oscillation periods found here but also on the order of weeks. This may explain the different periods of symptom fluctuations observed in previous studies (68). Future studies based on the reciprocal inhibition model may result in the development of new therapies and a deeper understanding of the pathogenesis of PTSD.

The current study has several limitations. First, we used avoidance symptoms as an index of the dissociative level despite this not being included in the official definition of the dissociative subtype. We used this measurement because the reciprocal inhibition model provides a prediction that the differences between PTSD subtypes are quantitative rather than qualitative. Unlike other dissociative symptoms, avoidance symptoms are found, to some extent, in all patients with PTSD. The reciprocal inhibition model predicts that oscillations between the two states should be observed in all patients with PTSD; however, the frequency and/or magnitude of these oscillations should differ between PTSD subtypes. Consistent with this, the findings of our behavioral experiment revealed that the relative frequency of the non-dissociative state, as indexed by the number of toward trials, was numerically but not statistically greater in patients with the non-dissociative compared with those with the dissociative subtype (t = 1.38, df = 18, *p* = 0.18). However, future studies should confirm this because the medium effect size (Hedge’s d = 0.60) indicates that a sample size of n = 44 for each group is required to reach statistical significance. Despite it is yet to empirical examination, this might have been a result of the average suppression of the amygdala by the vmPFC being relatively stronger in the dissociative subtype and vice versa in the non-dissociative subtype.

Second, there was non-uniformity in the studies included in the meta-regression analysis regarding several factors such as the preprocessing methods or experimental conditions. However, despite these methodology differences, we still observed a strong correlation between symptom imbalance and amygdala activity, which indicates the robustness of our findings.

Third, we only selected 9 studies for the meta-regression analysis. However, each study reported the results for dozens of people (total N = 316).

Fourth, although neural evidence, such as data from functional magnetic resonance imaging (fMRI), may support individual fluctuations and a pivotal role of vmPFC, we did not obtain this information in the current study. Therefore, there is a need for future neuroimaging studies on patients with PTSD focusing on within-individual dynamics. Although much more extensive and intensive examinations of our reciprocal inhibition model are necessary, we believe that the proposed model could be useful in providing a new perspective on the advancement of the diagnosis and treatment of PTSD. Classifying patients with PTSD with different symptoms into different subtypes has allowed more careful analysis of their differential responses to psychological trauma, which is eventually expected to lead to a more sophisticated understanding of the neurobiology and treatment of PTSD (69). The next step could be classification of different PTSD states within individual patients.

Overall, our reciprocal inhibition model may explain the associations between PTSD symptoms, attentional biases, and neural states. Our results are consistent with the concept that the neural state of a patient with PTSD may not be stable. Instead, individuals may fluctuate between dissociative and non-dissociative states. The reciprocal inhibition model proposed here may be used as a unifying framework to study the dynamics of diverse characteristics of PTSD, such as neural activities, attentional biases, and symptomatology.

## Acknowledgments

We thank K. Nakamura and C. Hosomi for their help with scheduling and conducting the experiment, A. Cortese, V. Taschereau-Dumouchel, S. Morinobu, and I. Liberzon for comments on the manuscript, and H. Moriya and A. Cortese for discussions about the experiments. The study was conducted in the ImPACT Program of Council for Science, Technology and Innovation (Cabinet Office, Government of Japan). This research was also supported by AMED under Grant Number JP18dm0307008.

## Author contributions

T.C., K.I., M.K., and A.K designed the study; T.C., K.I., S.K., and M.S. performed research experiments; T.C. analyzed data; and T.C., K.I., S.B., J.T., H.T., T.K., S.K., Y.H., A.H., T.M., T.Y., M.S., I.S., M.K., and A.K. interpreted the data and wrote the paper.

