## Supplementary Materials for "A reciprocal inhibition model of alternations between non-dissociative and dissociative states in patients with PTSD"

**Online-Only Supplements**

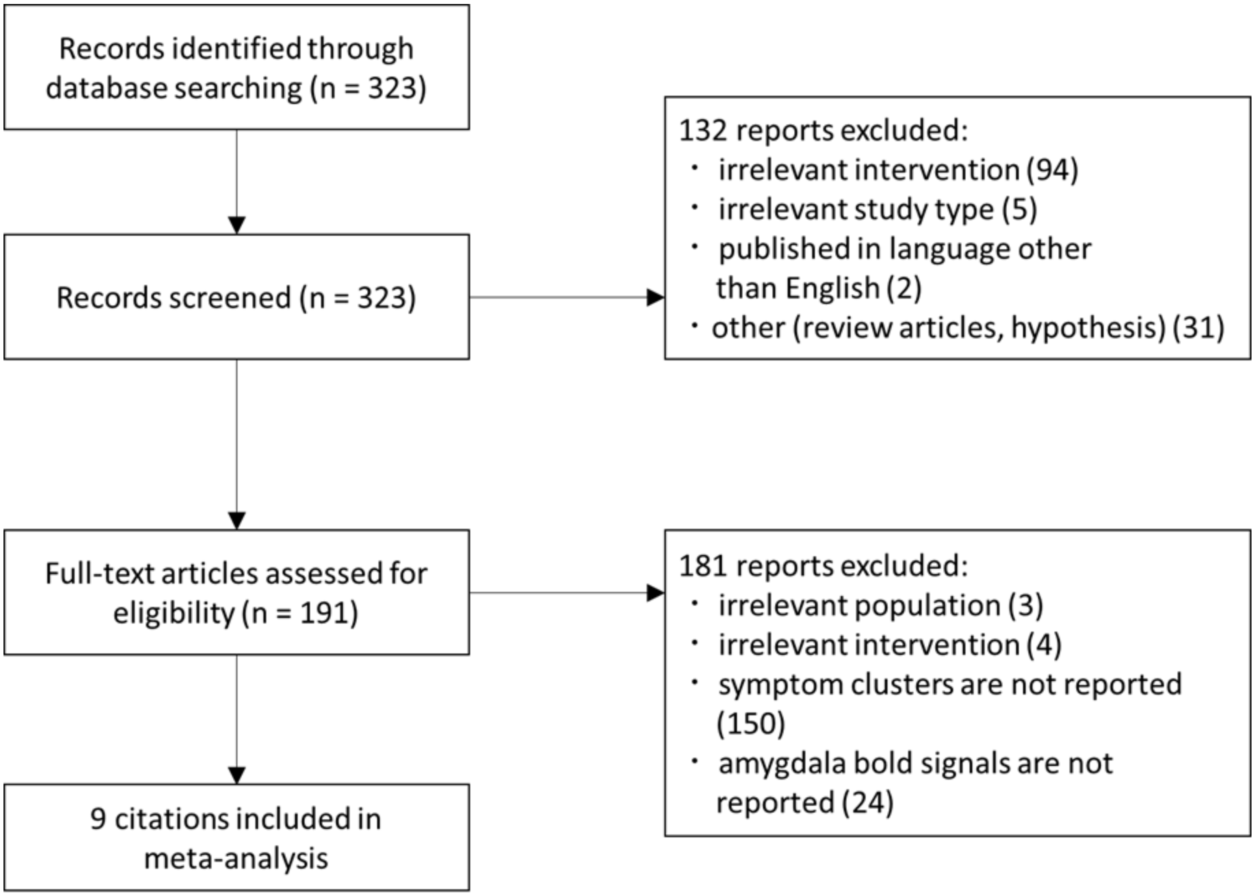


**eFigure 1. PRISMA chart for study selection.**

### eTable 1. Demographic data of PTSD patients

| ID | Sex | Age | Trauma type | Re-experiencing | Avoidance | Hypervigilance | Dissociation | Dissociative subtype* | AB TOWARD | AB AWAY |
|---|---|---|---|---|---|---|---|---|---|---|
| 1 | Female | 38 | Unpleasant sexual experience | 16 | 36 | 20 | 0 | 0 | 0.64 | 1.14 |
| 2 | Female | 48 | Domestic violence | 11 | 28 | 23 | 0 | 0 | 0.46 | 0.45 |
| 3 | Female | 53 | Domestic violence, Childhood abuse, Unpleasant sexual experience | 27 | 46 | 23 | 0 | 0 | 0.59 | 1.52 |
| 4 | Female | 46 | Domestic violence, Childhood abuse, Unpleasant sexual experience | 35 | 48 | 36 | 0 | 0 | 1.37 | 1.45 |
| 5 | Female | 51 | Domestic violence | 31 | 45 | 34 | 0 | 0 | 1.32 | 1.78 |
| 6 | Female | 38 | Domestic violence, Unpleasant sexual experience | 13 | 18 | 20 | 0 | 0 | 0.80 | 0.90 |
| 7 | Male | 53 | Childhood abuse, Unpleasant sexual experience | 26 | 39 | 11 | 0 | 0 | 1.46 | 1.39 |
| 8 | Female | 35 | Childhood abuse, Unpleasant sexual experience | 15 | 22 | 17 | 2 | 0 | 1.28 | 1.34 |
| 9 | Female | 53 | Domestic violence | 16 | 34 | 31 | 0 | 0 | 0.73 | 1.25 |

| ID | Sex | Age | Trauma type | Re-experiencing | Avoidance | Hypervigilance | Dissociation | Dissociative subtype* | AB TOWARD | AB AWAY |
|---|---|---|---|---|---|---|---|---|---|---|
| 10 | Female | 29 | Domestic violence, Childhood abuse | 24 | 23 | 26 | 0 | 0 | 0.59 | 0.46 |
| 11 | Female | 24 | Domestic violence, Childhood abuse | 22 | 37 | 27 | 3 | 1 | 0.55 | 0.43 |
| 12 | Female | 31 | Domestic violence, Unpleasant sexual experience | 14 | 30 | 14 | 8 | 1 | 0.58 | 0.81 |
| 13 | Female | 22 | Unpleasant sexual experience | 14 | 24 | 25 | 6 | 1 | 0.87 | 0.93 |
| 14 | Female | 41 | Domestic violence, Unpleasant sexual experience | 34 | 47 | 30 | 12 | 1 | 1.52 | 1.43 |
| 15 | Female | 49 | Childhood abuse, Unpleasant sexual experience | 23 | 37 | 26 | 3 | 1 | 0.18 | 0.26 |
| 16 | Female | 47 | Domestic violence | 16 | 25 | 18 | 5 | 1 | 0.34 | 0.80 |
| 17 | Male | 29 | Childhood abuse | 26 | 31 | 25 | 4 | 1 | 0.58 | 0.64 |
| 18 | Female | 46 | Domestic violence, Childhood abuse, Unpleasant sexual experience | 25 | 38 | 24 | 3 | 1 | 1.57 | 1.97 |
| 19 | Female | 38 | Childhood abuse | 27 | 45 | 36 | 8 | 1 | 0.34 | 0.81 |

| ID | Sex | Age | Trauma type | Re-experiencing | Avoidance | Hypervigilance | Dissociation | Dissociative subtype* | AB TOWARD | AB AWAY |
|---|---|---|---|---|---|---|---|---|---|---|
| 20 | Female | 53 | Domestic violence | 31 | 24 | 23 | 7 | 1 | 0.95 | 0.81 |
| Mean | - | 41.2 | - | 22.3 | 33.85 | 24.45 | 3.05 | - | 0.84 | 1.03 |

Symptoms are measured by the Clinician-Administered PTSD Scale for DSM–4.
*In order to meet the criteria for the dissociative subtype, individuals must additionally report as having depersonalization and/or derealization. We used the “1-2 rule” to define the presence of these symptoms, counting a symptom as present if it occurred at least monthly with at least moderate intensity.

### eTable 2. Summary of the characteristics of the studies included in the meta-analysis

| Author | Scale | Condition | Populations (N) | Imbalance | Re-experiencing* | Avoidance* | ltAmyg** | rtAmy** |
|---|---|---|---|---|---|---|---|---|
| Naegeli[1] | CAPS | white noise burst > baseline | PTSD (28) | 0.016 | 0.463 | 0.446 | 4.54 | N.S. |
| Lieberman[2] | SCID*** | happy, angry, fearful, sad faces > shapes | TEI (45) | 0.033 | 0.298 | 0.265 | 5.57 | 5.98 |
| Stevens[3] | PSS | fear > neutral face | PTSD+TEI (31) | 0.018 | 0.313 | 0.295 | N.S. | 3.44 |
| Frijling[4] | CAPS | fear > happy face | ASD (41) | 0.237 | 0.423 | 0.186 | N.S. | 4.28 |
| Rabinak (2014)[5] | CAPS | unpleasant > neutral images | PTSD (21) | -0.008 | 0.420 | 0.429 | 3.23 | 4.60 |
| | | | TEI (21) | -0.020 | 0.010 | 0.030 | 3.65 | 3.37 |
| Stevens (2014)****[6] | PSS | fear > neutral face | PTSD+TEI (22) | -0.015 | 0.247 | 0.262 | 3.52 | 3.12 |
| vanRooij (2014)[7] | CAPS | unpleasant > neutral images | TEI (28) | 0.001 | 0.018 | 0.017 | 4.76 | 6.00 |
| | | | HC (25) | -0.004 | 0.017 | 0.021 | N.S. | 5.65 |
| Aupperle (2013)[8] | CAPS | anticipation of unpleasant > neutral images | PTSD (14) | 0.073 | 0.520 | 0.446 | N.S. | 2.96 |
| Stevens (2013)****[9] | PSS | fear > neutral face | PTSD (20) | -0.071 | 0.467 | 0.538 | 3.13 | 2.69 |
| | | | TEI (20) | -0.008 | 0.107 | 0.114 | 2.65 | 2.31 |

TEI: trauma-exposed individuals, ASD: acute stress disorder, HC: healthy control
*PTSD symptoms were normalized such that the highest possible value was one and the lowest possible value was zero.
**Amygdala activations are described as z-scores.
***The PTSD scale was measured using the structured clinical interview for DSM-V (SCID). Avoidance and numbing symptoms were combined into one category to allow for direct comparison of studies that used the DSM-IV with those that used the DSM-V.
****The subsets of populations from these two studies are partially overlapping.
